# Neophobic response of bull ants (*Myrmecia midas*) to odours introduced on their foraging route

**DOI:** 10.1101/2024.07.15.603486

**Authors:** Venkata Manvitha Kambala, Yousef Ahmed, Jasmine Lee, Anwin Jose, Sahera Syed Nadir, B C Priyanka, Ali Gabir, Yingdie Sun, Ken Cheng, Sudhakar Deeti

**Affiliations:** School of Natural Sciences, Macquarie University, Sydney, NSW 2109, Australia

**Keywords:** Sensory Cues, Eusociality, Detour, Chemical Cues, Insect navigation

## Abstract

Goal-oriented learning and navigation is well known in eusocial insects. The solitary foraging of nocturnal bull ants *Myrmecia midas* in their visually complex environment relies on path integration and landmark learning. While this species seems to be ‘sensitive’ to handling and reacts to visual changes in their surroundings, not much is known about how added olfactory stimuli impact their route navigation on a vertical surface. In the current study, we added one of five different invisible odours on the trees on which foragers normally forage. We found that the bull ants showed neophobic responses to all the odours. The Tea-tree and Lavender odours showed the strongest impact on the bull ants’ navigation by causing detours, U-turns, and avoidance of the sensory stimuli, with the ants meandering more and scanning more frequently. The odours of Olive oil, Flax-seed oil, and Eucalyptus oil had a moderate impact on the ants’ navigation. These findings showed the widespread influence of non-visual chemical cues in shaping bull ant navigation and highlight the induction of neophobic responses stemming from chemical alterations on learned routes. Overall, this study contributes to the understanding of the effects of foreign odours, adding to our understanding of the complex learning processes of bull ants in their vertical navigation.

## INTRODUCTION

Insects possess small brains with limited processing power, yet exhibit sophisticated and complex behaviours. One extraordinary feature is the ability to efficiently navigate from home to their destination using a navigational tool kit consisting of multiple strategies (Buehlmann et al., 2020; Wehner, 2020). Ant species that conduct group-based foraging rely on conspecific cues, typically pheromones, to navigate (Knaden and Grahan, 2016). A considerable number of ant species, however, forage solitarily, whereby each individual ant must complete its own search for food and return to its nest. These solitary foragers are adept at navigating on the ground, on trees, or underground. Bull ants, for example, are skilled in recognising their environments, the route to a foraging tree to locate food sources, and the way back to their nest (Freas et al., 2017; Narendra et al., 2013).Navigating ants use multiple strategies (Wehner, 2008, 2020), including path integration (PI) (Collett and Collett, 2000; Wehner and Srinivasan, 2003) and learned panorama cues (Collett et al., 2006; Collett, 2012; Freas et al. 2019; Schultheiss et al., 2016; Wehner, 2003; Wystrach et al. 2011; Zeil and Fleischmann 2019). Visual panoramas are thought to be learned by performing learning walks, widely observed behaviours in desert ant genera such as *Cataglyphis* (Fleischmann et al., 2016; Wehner et al., 2004), *Ocymyrmex* (Müller and Wehner, 2010), and *Melophorus* (Deeti and Cheng, 2021; Deeti et al., 2024a). Laerning walks are not exclusive to arid environments, as they are also exhibited by non-desert ant species such as *Myrmecia* bull ants (Jayatilaka et al., 2018) and *Formica* wood ants (Nicholson et al., 1999). Ants are also known to use olfactory cues for locating food sources (Wolf and Wehner, 2000) and their nest (Steck et al., 2009) and marking trails to guide nestmates to resources (Buehlmann et al., 2020; Knaden and Graham, 2016). The prevalence of these behaviours in diverse ant species highlights the importance of olfactory cues in navigation, foraging, and place learning. However, the importance of olfactory cues on bull ants’ vertical navigation is largely unknown.

The bull ant species *Myrmecia midas* inhabits underground structured nests and journeys up surrounding trees to forage for food, then successfully descending down the tree to return to the nest (Reid et al., 2011; Narendra et al., 2013; Islam et al., 2021, 2022). Nocturnal members of this genus, including *M. pyriformis* and *M. midas*, have an added challenge: conducting these three-dimensional foraging trips during evening and morning twilight when low light levels make visual cues harder to detect. Ants are known to use landmarks to navigate, and they can learn to associate landmarks with food sources (Collett and Collett, 2002). *M. midas* navigates visually while descending its foraging tree (Freas et al., 2018). Occasionally, while descending the tree trunk, *M. midas* foragers scan by bringing their head and eyes to a near upright position while their body remains vertically oriented. This behaviour suggests that whilst descending, *M. midas* foragers use visual cues memorised while horizontal on the ground to steer to the correct side of the tree. Blocking the panoramic views surrounding the tree leads to disorientation in the descent. Ascending foragers climb up the tree trunk to the canopy, but have not been observed to make these scans (personal observations), suggesting that they may attend to different cues, possibly ‘vertical views’ of the canopy, to orient while climbing. Visual changes, such as movement or sudden appearance or disappearance of surrounding landmarks, can prompt these ants to react because bull ants, like many other ants, rely heavily on their vision to navigate their environment and find food (Collett et al., 2006; Islam et al., 2021, 2022, 2023). Ants have compound eyes that allow them to detect even the slightest movements and changes to their surroundings (Warrant and Dacke, 2010, 2011).

Novel odours on an animal’s foraging path might generate a phobic response. Neophobia leads to an avoidance of unfamiliar stimuli, serving as a protection mechanism from possible threats heralded by the novel stimuli. The classification of novel and unfamiliar applies to anything that an organism has not experienced in the context in which the stimulus appears (Greenberg and Mettke-Hofmann, 2001). Novelty can be found in food, circumstances, environments, fluids, and objects. Whilst neophobia varies across species, in birds, it increases with the degree of unfamiliarity (Miller et al., 2022). Neophobic responses may be reduced if an animal is exposed to a specific stimulus repeatedly (Greenberg and Mettke-Hofmann, 2001). Bull ants *M. midas* exhibit neophobic responses to changes in their learned routes (Islam et al., 2023). This includes detouring around new or unfamiliar objects, meandering more, and scanning the environment more frequently when presented with tactile and olfactory changes on the tree on which they travel to forage for food. Here we further explore their response to olfactory changes on their learned route with a range of chemical cues, examining changes in their travelling behaviour in more detail.

In this study, we explored how the ascending navigational behaviour of nocturnal bull ant *Myrmecia midas* foragers is influenced by introduced olfactory cues placed on their established foraging routes. Firstly, we made control or baseline observations on the travel characteristics of bull ants when travelling vertically up the eucalyptus tree trunk. Then on the following day, we examined possible neophobic responses of the bull ants by applying a new olfactory cue on their foraging tree. The novel odours consisted of Eucalyptus oil, Olive oil, cold-pressed Flax-seed oil, Tea-tree oil, and Lavender oil. The direct route required walking over the added odours. We hypothesised that by introducing novel odours on their learned route, we would trigger detour or avoidance behaviour among the foragers. To assess the impact of the foreign odours, we measured path characteristics such as sinuosity and path straightness, speed, angular velocity of orientation shifts, trajectory duration, the number of scanning bouts, and the scanning bout durations, comparing control and test (odour-added) conditions. The behavioural changes will allow us to understand how neophobic responses of the bull ants to the chemical cues manifest in behaviour.

## MATERIALS AND METHODS

### Study location

During the Australian autumn months from March to April 2024, we conducted odour-response studies on nocturnal bull ant *Myrmecia midas* colonies at the Macquarie University campus, North Ryde, Sydney, Australia (33°46′18″ S, 151°06′30″ E). These bull ants typically construct their ground-nesting colonies near the base of eucalyptus trees, within a range of about 30 cm, to gain elevation and safeguard against floods (Deeti et al., 2024b). From our observations, we noticed that ants begin foraging vertically on the eucalyptus trees during the evening twilight, immediately after sunset, and their activity peaks within 20 to 30 minutes. They then return to the nest during the morning twilight. These experiments were conducted exclusively on trees near the nest, specifically focusing on foragers on the nest tree. Although ethical approval was not required for researching and collecting ants in Australia, we ensured that our experiments did not harm individual ants or their colonies.

### Experimental setup

We selected five different trees and nests for this experiment. Before conducting the experiments, we observed the individual trees for a week to understand the ants’ well-maintained foraging corridor while vertically travelling up on the tree, and then we marked their preferred foraging corridor. All the selected foragers had followed a 40-cm-wide corridor. Before conducting any experiments, we labelled the four corners of the recording area, situated 50 cm above the ground, to mark a 1-m-high and 70-cm-wide area where we could record the paths of individual foragers during their upward foraging trips, which we call ascending navigation, distinct from their ground-level movements. We used a Sony Handy camera (FDR-AX700) to record the paths of vertically ascending navigators. The camera was set on a tripod positioned 0.6 m above the ground, recording footage at 25 frames per second. The camera’s field of view covered the focal area of the tree surface, recording with a high resolution of 3860 by 2160 pixels. The camera was turned on before the first forager entered the recording area and continuously recorded for 40 minutes.

### Procedure

In our experiments, we used five different odours for investigation. These included Eucalyptus oil (Coles^®^, Eucalyptus Oil 1 mL/mL), representing a local odour of the experimental tree surface, Olive oil (Remano^®^, oleic acid 0.75 mL/mL), which is a source rich in oleic acid, and cold-pressed Flax-seed oil (Coles^®^, linoleic acid, 0.63 mL/mL), a source rich in linoleic acid, cold-pressed to preserve this fragile omega-6 fatty acid. Both these fatty acids have been recognized as odours that ants associate with death (Sun et al. 2018). We also included Tea tree oil (Thursday Plantation^®^, Melaleuca Oil 1 mL/mL) and Lavender oil (Hemani^®^, Lavender oil 1 mL/mL) as foreign odours in our study. All these odours were novel on the foraging route. Each of these odours was tested only once on the nest tree of a single nest, different nests for different odours. The experiment at each nest lasted two days, with Control observations on day 1 and Test-day observations on day 2. We did not capture or paint any ant because this process is aversive to this species and will generate avoidance responses whether any odour is added or not (Lionetti et al., 2023).

### Control day

On the control recording day, we set up the camera 5 minutes before sunset. Once a forager reached the base of the foraging tree, the observer turned on the camera to record. The recording continued for 40 minutes or until the sample size reached 15 ants. We did not apply any scents on the control day.

### Test day

On the test day, 5 minutes before sunset, we placed a scent in the middle of the recording area along the foraging path. We used 2 ml of oil on each tree, a horizontal strip 50 cm by 5 cm applied with a cotton gauze. Once the scent was applied, the observer began recording with the camera as soon as a forager ant reached the base of the foraging tree. The recording continued for the next 40 minutes or until 15 ants had been observed.

### Tracking

We used the animal tracking program DLTdv8 (version 8.2.9) in MATLAB (2022B) to extract frame-by-frame coordinates of the head and thorax — specifically the tip of the head and the middle of the thorax (see Fig. S1) — for each ant in every video obtained during our recording of control and test days. These extracted frame-by-frame coordinates served as the basis for all subsequent analyses of the foragers’ movements and behaviour.

### Data analysis

To understand how quickly and far the ants moved, we calculated speed. Speed refers to the magnitude of an ant’s velocity and was calculated as the average over the entire videotaped trajectory for each ant, excluding the stopping durations. We also calculated the orientation angular velocity by measuring the rate of change of an ant’s orientation direction over time. When walking, the orientation direction of these ants typically oscillates continuously, with the head swinging left and right. Orientation direction in any frame was determined as the direction of the straight line passing from the thorax coordinates through the head coordinates. Orientation angular velocity was calculated by dividing the change in orientation angle by the corresponding change in time, providing a measure of how quickly the ant was altering its head direction. To understand path characteristics, we used three indices of straightness: *path straightness*, *sinuosity*, and *E ^a^_max_*, each of which relates to the directness of navigation away from the nest. Path straightness is a measure that quantifies the extent to which an ant’s path deviates from a straight line between its starting point on the video and its endpoint, reflecting the directness of navigation. It is defined as the straight-line distance of the path on video divided by the path length. Straightness ranges from 0 to 1, with larger values indicating straighter paths, while smaller values indicate more curved or convoluted paths. Sinuosity assesses the degree of waviness or curvature in an ant’s path as it navigates away from the nest (Batschelet 1981; Deeti et al. 2023a, 2024a). A higher sinuosity value indicates a more convoluted path, while a lower value indicates a straighter path. *Sinuosity* is an estimate of the tortuosity in a path, calculated as 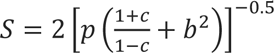, where *p* is the mean step length, *c* is the mean cosine of turning angles and *b* is the coefficient of variation of the step length. A trajectory *step* is the movement between the positions of the animal (thorax positions) recorded at consecutive video frames. Accordingly, step lengths are the Euclidean distances between consecutive points along a path, and turning angle refers to the change in direction between two consecutive steps (Benhamou, 2004). The maximum expected displacement of a path, *E ^a^ _max_* = β/(1–β), where β is the mean cosine of turning angles, is a dimensionless value expressed as a function of number of steps, and is consistent with the intuitive meaning of straightness (Cheung et al., 2007). Larger maximum expected displacement values indicate straighter paths, hence greater displacement, while a smaller value suggests more localised or constrained movement.

During the outbound navigation, ants frequently displayed a series of stereotypical successive fixations in different head directions by stopping and rotating on the spot at one location, known as a “scanning bout” (Deeti et al., 2023b). During each ant’s control and test run, we extracted the number of scanning bouts and the scanning-bout durations, from the start of a scanning bout until the ant started walking again.

### Statistical analysis

The experiments conducted at the five different nests were analysed using t-tests, one test for each nest.

## RESULTS

After emerging from the nest during the evening twilight, all the nest-tree foragers followed a stereotypical vertical path up the eucalyptus tree trunk and continued along the same foraging corridor every day. We conducted observations on these experienced foragers for two consecutive nights on their nest trees for control and odour-test purposes. The presence of odours on their foraging corridor resulted in detours for foraging (Fig. 1). Specifically, with Eucalyptus oil, 46.4% of ants walked over the odour, while 53.6% took detours to avoid it. With Olive oil and Flax-seed oil (sources of oleic acid and linoleic acid, respectively), 35% and 40% of foragers walked over the odour, while 65% and 60% took detours. With Tea tree oil and Lavender oil, none of the foragers walked over the odour, with 26% and 22% of foragers taking a U-turn and returning to the nest, while 74% and 78% took detours away from the odour line. These results suggest that foragers exhibit a neophobic response, with nearly half of them avoiding the familiar Eucalyptus oil, more avoiding oils containing oleic and linoleic acid, and complete avoidance of Tea tree and Lavender oils.

**Fig. 1.**
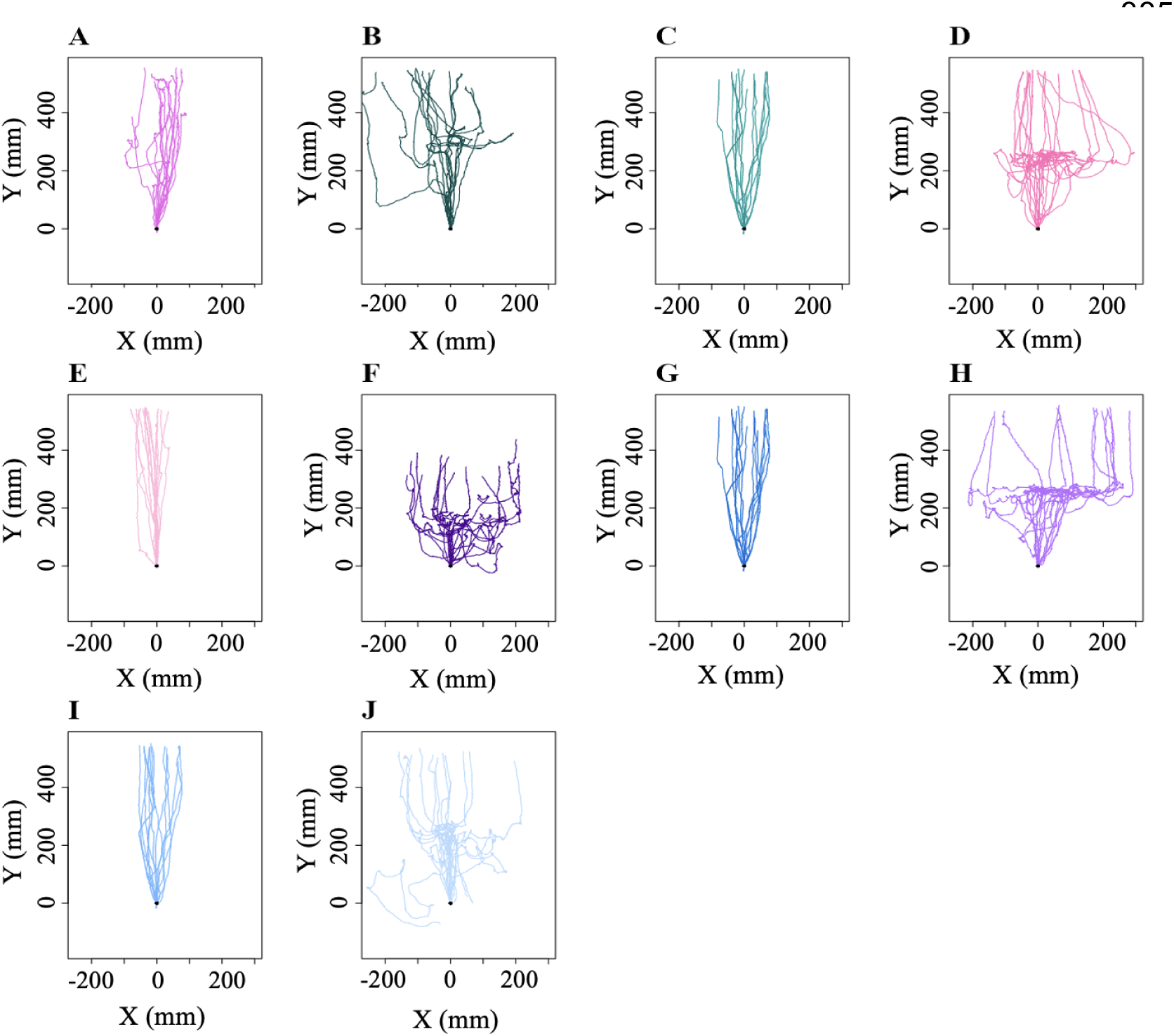
Trajectories of ants in control and test conditions with various odours added. (A) to (J) show the Control and Test paths of the bull ants during vertical navigation. (A), (C), (E), (G), and (I) represent the control paths before Eucalyptus oil, Olive oil, cold-pressed Flax-seed oil, Tea tree oil, and Lavender oil, respectively, were added. (B), (D), (F), (H), and (J) depict the test paths after Eucalyptus oil, Olive oil, cold-pressed flax-seed oil, Tea tree oil, and Lavender oil, respectively, were added. Each trajectory plot represents the movement path of an individual ant. The y-axis points upwards. The coordinates (0,0) represent the starting frame of each recording.

In path characteristics, ants exhibited a more curved and meandering trajectory in the odour-change conditions compared to the control conditions. We found differences across conditions in each of our three measures of path meander. Firstly, sinuosity increased significantly in the odour change condition (Olive oil control vs. Olive oil test: *t* = –7.47, *df* = 17.41, *P* < 0.0001; Flax-seed oil control vs. Flax-seed oil test: *t* = –14.12, *df* = 19.49, *P* < 0.0001; Tea tree oil control vs. Tea tree oil test: *t* = –10.41, *df* = 18.22, *P* < 0.0001; Lavender oil control vs. Lavender oil test: *t* = –12.2, *df* = 22.81, *P* < 0.00001) except for Eucalyptus oil (Eucalyptus oil control vs. Eucalyptus oil test: *t* = 0.4, *df* = 23.85, *P* = 0.68) (Fig. 2A). Secondly, E ^a^ _max_ was lower in the odour-change conditions, with the ants having a smaller amount of displacement per unit length travelled compared to the control conditions. The t-tests showed a significant difference in test conditions compared to the control conditions in all comparisions except Eucalyptus oil (Olive oil control vs. Olive oil test: *t* = 6.18, *df* = 25.23, *P* < 0.0001; Flax-seed oil control vs. Flax-seed oil test: *t* = 7.28, *df* = 16.86, *P* < 0.0001; Tea tree oil control vs. Tea tree oil test: *t* = 9.08, *df* = 16.98, *P* < 0.0001; Lavender oil control vs. Lavender oil test: *t* = 9.64, *df* = 23.23, *P* < 0.0001; Eucalyptus oil control vs. Eucalyptus oil test: *t* = –0.02, *df* = 27.59, *P* = 0.98) (Fig. 2B). With straightness, the odour change led to a significantly higher magnitude of path deviation from a straight-line path compared to paths in the Control conditions, again in all comparisons except Eucalyptus oil (Olive oil control vs. Olive oil test: *t* = 6.47, *df* = 14.27, *P* < 0.001; Flax-seed oil control vs. Flax-seed oil test : *t* = 10.2, *df* = 15.52, *P* < 0.0001; Tea tree oil control vs. Tea tree oil test : *t* = 8.64, *df* = 14.32, *P* < 0.0001; Lavender oil control vs. Lavender oil test: *t* = 5.03, *df* = 14.11, *P* < 0.001; Eucalyptus oil control vs. Eucalyptus oil test: *t* = 1.75, *df* = 26.13, *P* = 0.09) (Fig. 2C).

**Fig. 2.**
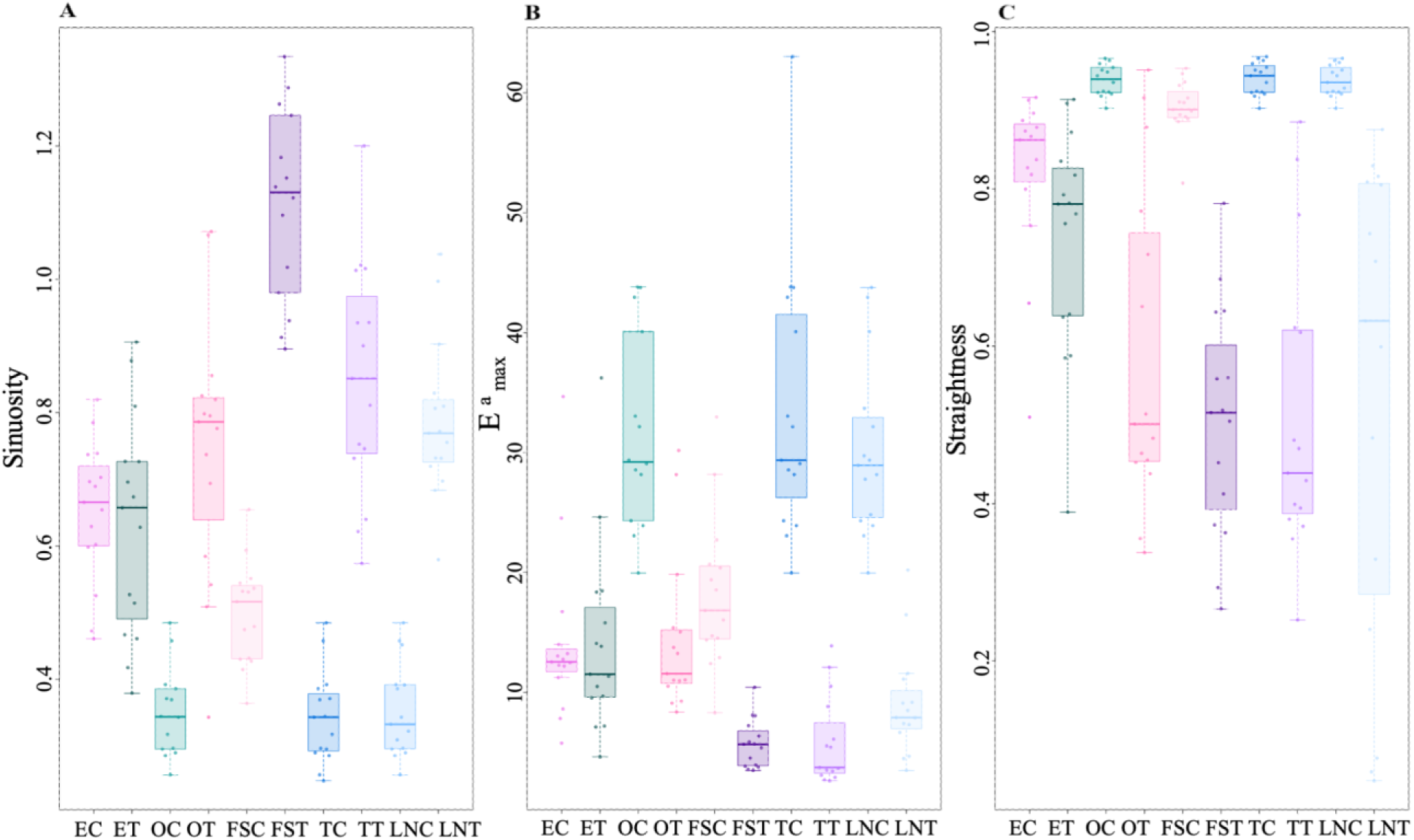
Path characteristics of bull ants during the control and test odour trials. Shown are the (A) Sinuosity, (B) E^a^, and (C) Straightness under control and test-odour observations. Box plots represent the median (line inside the box), interquartile range (box), and extreme values excluding outliers (whiskers). Individual data points are shown as dots. Each point represents a single trajectory measure. EC, OC, FSC, TC, LNC represent Eucalyptus oil, Olive oil, Flax-seed oil, Tea tree oil, and Lavender oil controls whereas ET, OT, FST, TT, LNT represent Eucalyptus oil, Olive oil, Flax-seed oil, Tea tree oil, and Lavender oil tests respectively.

Odour changes had a noticeable impact on the speed, orientation angular velocity, and duration of the foragers. When the foragers sensed non-visual odour stimuli, they slowed down, frequently shifted their gaze in different directions, and stayed in the recording area longer compared to the respective controls (Fig. 3). In speed, the t-tests revealed significant differences in all comparisons (Fig. 3A), with the ants speeding up with Eucalyptus oil and slowing down with other odours (Eucalyptus oil control vs. Eucalyptus oil test: *t* = –3.04, *df* = 27.63, *P* < 0.01; Olive oil control vs. Olive oil test: *t* = 6.57, *df* = 26.63, *P* < 0.0001; Flax-seed oil control vs. Flax-seed oil test: *t* = 5.21, *df* = 27.01, *P* < 0.0001; Tea tree oil control vs. Tea tree oil test: *t* = 8.54, *df* = 24.25, *P* < 0.0001; Lavender oil control vs. Lavender oil test: *t* = 13.94, d*f* = 19.06, *P* < 0.00001). Changes in odour also increased the magnitude of orientation angular velocity in the foragers (Fig. 3B). The t-tests showed significant differences between all pairs in orientation angular velocity except for Olive oil (Eucalyptus oil control vs. Eucalyptus oil test: *t* = –4.64, *df* = 15.29, *P* < 0.001; Flax-seed oil control vs. Flax-seed oil test: *t* = –20.84, *df* = 15.16, *P* < 0.00001; Tea tree oil control vs. Tea tree oil test: *t* = −8.94, *df* = 21.12, *P* < 0.0001; Lavender oil control vs. Lavender oil test: *t* = –9.06, *df* = 25.11, *P* < 0.0001; Olive oil control vs. Olive oil test: *t* = 0.33, *df* = 20.93, *P* = 0.73). The amount of time spent in the recording area also increased, as foragers stopped just before the odour line and meandered around it (Fig. 3C). Consequently, trips took significantly more time with all odours except for Eucalyptus oil (Olive oil control vs. Olive oil test: *t* = –5.3, *df* = 14.35, *P* < 0.0001; Flax-seed oil control vs. Flax-seed oil test: *t* = –4.8058, *df* = 16.4, *P* < 0.0001; Tea tree oil control vs. Tea tree oil test: *t* = –6.6, df = 14.66, *P* < 0.0001; Lavender oil control vs. Lavender oil test: *t* = – 5.1, df = 14.16, *P* < 0.0001; Eucalyptus oil control vs. Eucalyptus oil test: t = –0.70, *df* = 23.49, *P* = 0.48).

**Fig. 3.**
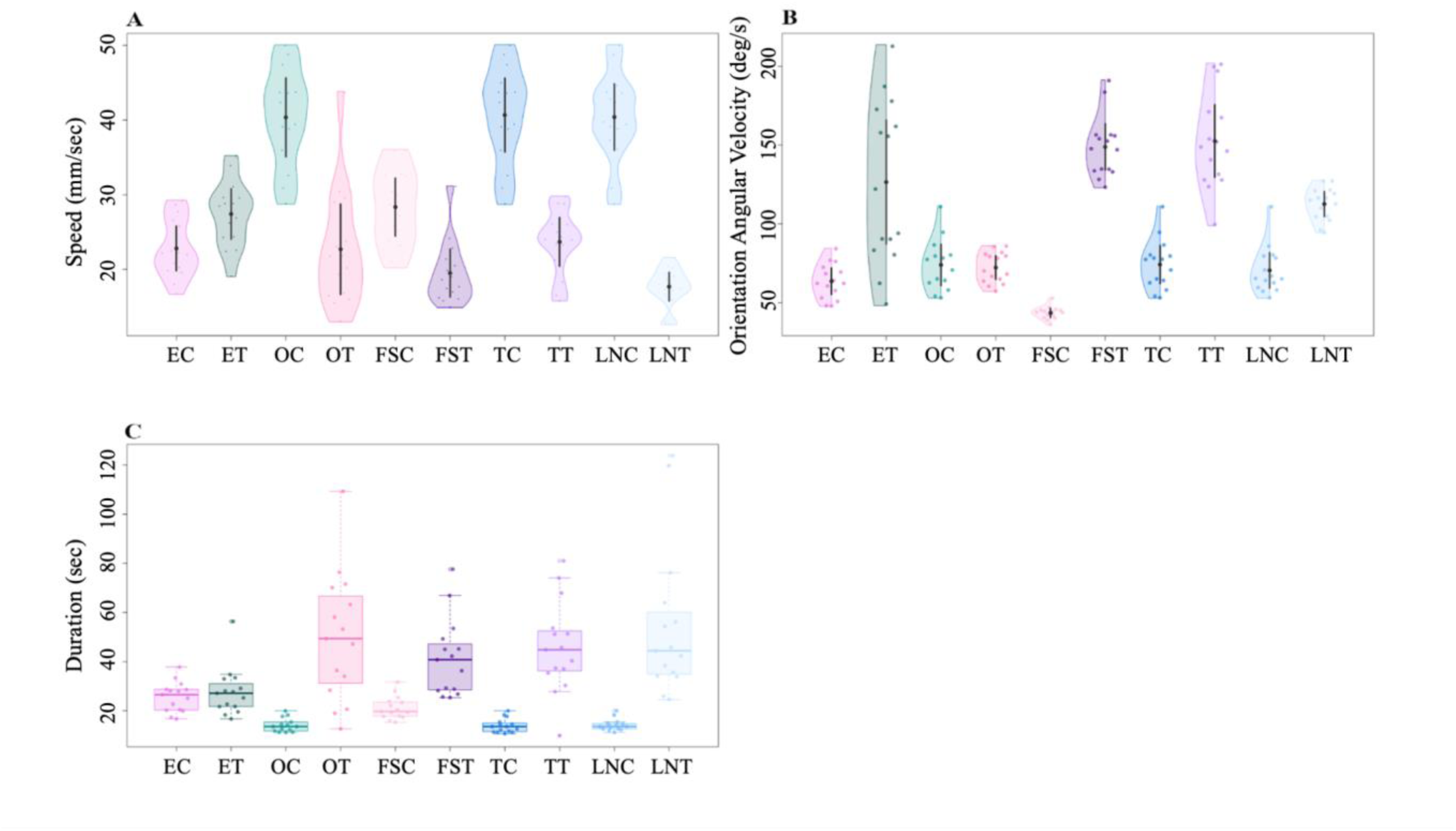
Comparison of Speed, Orientation Angular Velocity and Duration of the entire path of bull ants during the control and test odour trials. The violin (A) and half-violin (B) plots show the distributions of mean speed and orientation angular velocity of ants across control and test trials. In the violin and half-violin plots, the solid dot shows the mean while the vertical bar represents the 95% confidence interval of the mean. Box plots (C) represent the median (line inside the box), interquartile range (box), and extreme values excluding outliers (whiskers). Individual data points of the entire trajectories’ durations are shown as dots. Each point represents a single trajectory measure. EC, OC, FSC, TC, LNC represent Eucalyptus oil, Olive oil, Flax-seed oil, Tea tree oil, and Lavender oil controls and whereas ET, OT, FST, TT, LNT represent Eucalyptus oil, Olive oil, Flax-seed oil, Tea tree oil, and Lavender oil tests, respectively.

Odour change increased scanning in foragers. In Control recordings, i.e. those without any odour change, the majority of the foragers performed a single scan across their entire recording area (Fig. 4A), In contrast, on odour trials, where ants experienced a non-visual odour change, foragers scanned at least twice, with a maximum of 28 (Fig. 4A). The t-tests revealed significant differences in the number of scanning bouts in all comparisons (Eucalyptus oil control vs. Eucalyptus oil test: *t* = –4.79, *df*= 18.33, *P* < 0.001; Olive oil control vs. Olive oil test: *t* = –4.8, *df* = 15.17, *P* < 0.001; Flax-seed oil control vs. Flax-seed oil test: t = –4.10, *df* = 15.94, *P* < 0.001; Tea tree oil control vs. Tea tree oil test: *t* = –5.51, *df* = 14.93, *P* < 0.0001; Lavender oil control vs. Lavender oil test: *t* = – 4.72, *df* = 15.99, *P* < 0.001). The duration of scanning bouts also increased significantly with odour changes (Fig. 4B), in all comparisons except for Eucalyptus oil (Olive oil control vs. Olive oil test: *t* = –4.29, *df* = 14.47, *P* < 0.001; Flax-seed oil control vs. Flax-seed oil test: *t* = –2.77, *df* = 27.87, *P* < 0.01; Tea tree oil control vs. Tea tree oil test: *t* = –4.88, *df* = 14.56, *P* < 0.001; Lavender oil control vs. Lavender oil test: *t* = –5.42, *df* = 24.11, *P* < 0.0001; Eucalyptus oil control vs. Eucalyptus oil test: *t* = –1.13, *df* = 17.60, *P* = 0.27).

**Fig. 4.**
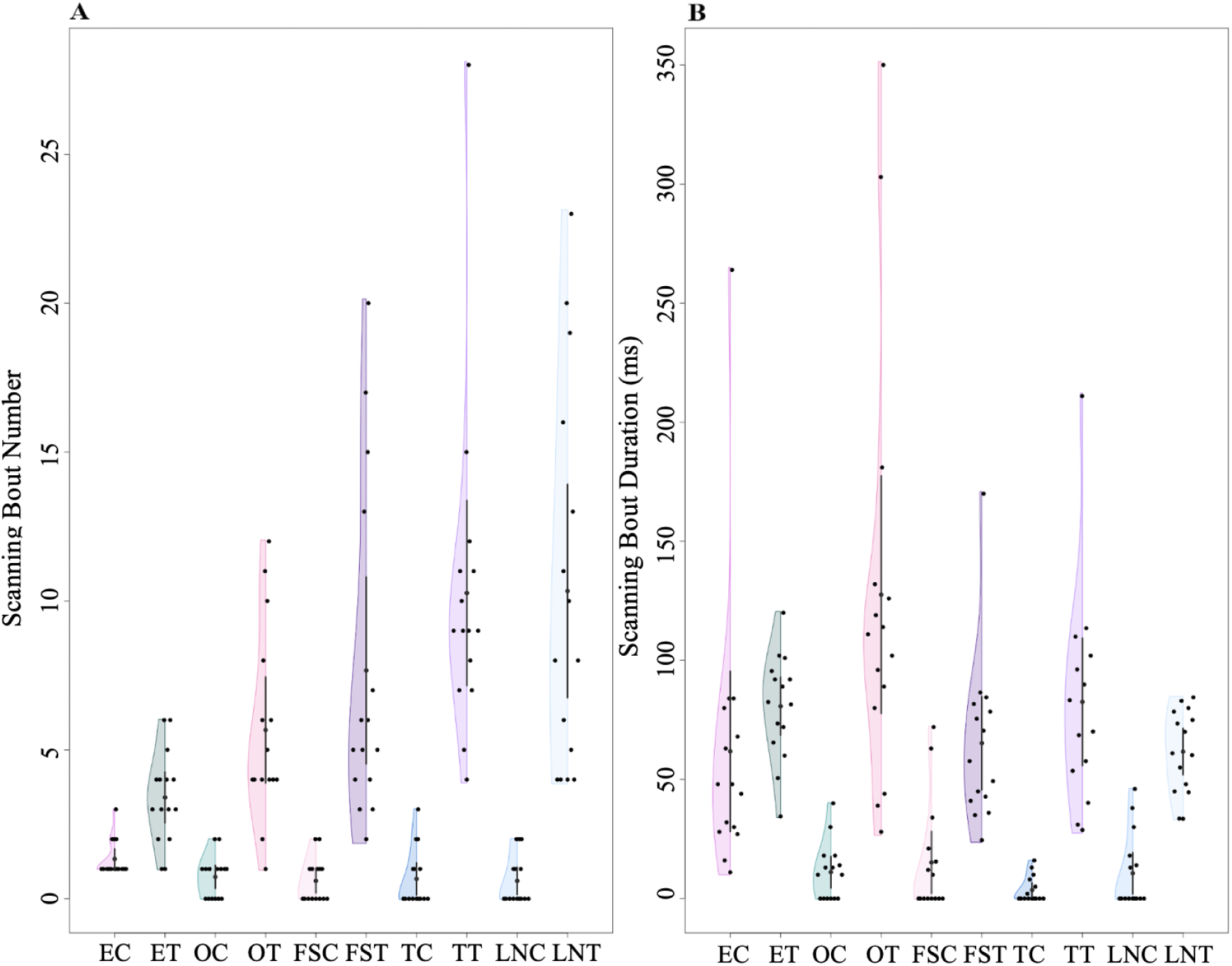
Number of scanning bouts and the duration of scanning bouts of the bull ants during the control and test observations. The half-violin plots in (A) and (B) show the distribution of the mean number of scanning bouts and duration of the scanning bouts, respectively, of ants across control and test trials. In the violin and half-violin plots, the solid dot shows the mean while the vertical bar represents the 95% confidence interval of the mean. EC, OC, FSC, TC, LNC represent Eucalyptus oil, Olive oil, Flax-seed oil, Tea tree oil, and Lavender oil controls and whereas ET, OT, FST, TT, LNT represent Eucalyptus oil, Olive oil, Flax-seed oil, Tea tree oil, and Lavender oil tests, respectively.

## DISCUSSION

Overall, this experiment examined whether manipulating bull ants’ natural environment with odourific oils will elicit abnormal behaviour. When exposed to foreign odours along their regular foraging path on the tree trunk, the ants immediately showcased neophobic behaviours, consistent with a previous study (Islam et al., 2023). Foragers exhibited avoidance behaviour. The level of avoidance varied depending on the odour, with complete avoidance for Tea tree and Lavender oils, moderate avoidance for oils containing oleic and linoleic acid, and partial avoidance for Eucalyptus oil, whose odour is familiar to them, albeit much stronger than usual. The results suggest that ants are most habituated to Eucalyptus oil, showcasing the least disturbance in their daily routine. None of the foragers walked over the Tea tree oil and Lavender oil; on average 76% detoured around the odour corridor, exhibiting the most neophobic response. In the presence of novel odours, the ants’ paths became more curved and meandering, as indicated by increased sinuosity, decreased straightness, and lower displacement efficiency compared to the control conditions. When encountering novel odours, the ants slowed down their speed, exhibited increased head-turning angular velocity (except for Olive oil), and spent more time in the recording area, suggesting they were searching for alternative routes. In the presence of odour changes, the ants performed multiple scanning bouts, with some individuals scanning up to 28 times, in contrast to a single scan in the control conditions. The duration of scanning bouts also increased significantly with odour changes. In summary, the study demonstrates that foraging ants exhibit neophobic responses when exposed to unfamiliar odours along their regular foraging paths. This response manifests as avoidance behaviour, path deviations, reduced speed, increased scanning, and altered movement patterns as they attempt to find alternative routes around the novel odour stimuli.

Animals are known to exhibit neophobic responses to changes in their learned routes by detouring around new stimuli or unfamiliar objects, meandering more, and scanning the environment more frequently (Mettke-Hofmann, 2022; Islam et al., 2023). The current study species, the nocturnal bull ant *M. midas*, showed a neophobic response when they were captured and released back at the same location after a minute (Lionetti et al., 2023) and the location became aversive to them. Neophobia was also obwrved in the context of odour changes introduced in their regular foraging pathways (Islam et al., 2023): altering the tactile or olfactory environment induced neophobic responses in the bull ants. Similarly, in our observations, changes in the odour on their foraging paths triggered neophobic behaviour. Here we found the extent of neophobic response to particular stimuli varied. For known odours such as Eucalyptus, Flax-seed, and Olive oil, the ants showed a moderate neophobic response. However, the foreign Tea tree and Lavender oils induced a stronger neophobic response. None of the foragers walked over these odours; instead, some took U-turns to return to their nest and all the ants’ paths were found to be more curved and less straight, with frequent stops and scans, likely due to their search for an alternative way to reach the tree canopy by taking detours from the original path. As a result, it took them longer to reach the final heading direction. We conjecture that the longer working life of this species and the smaller number of foragers in the colony are functional reasons for exhibiting a neophobic response (Lionetti et al., 2023). A longer working life and a smaller work force mean that each worker is more valuable to the colony. A more valuable life should be more cautious about changes in the environment, which might signal danger. Mechanistically, the ants likely perceived the non-visual unfamiliar odours as potentially harmful, causing them to avoid walking over them. Future studies should explore how these olfactory changes affect their foraging efficiency and behaviour over longer periods.

Our experiments found that non-visual odour changes on their newly learned route initiated detour learning in bull ants to avoid the odour line. Might the ants learn to detour more efficiently over trials? Detouring is utilised for effective navigation in a diverse range of species, including bull ants, in situations where they are required to adapt to unfamiliar physical stimuli and overcome barriers in their environments (Islam et al., 2023; Kabadayi et al., 2018; Zucca et al., 2005). In *M. midas*, setting up a barrier on the tree provided insights into their spatial detour learning (Islam et al., 2023). The nocturnal bull ants initially struggle to overcome the physical barrier, but, over multiple trials, they devised more efficient pathways, displaying their spatial learning abilities. They reshaped their route for more efficient navigation. These findings reveal the importance of learning to cope with changes in the surroundings and suggest that the detouring ants in our study would have detoured more efficiently were the experiment to carry on over multiple nights. Similar detouring behaviours were found in other species such as horses (Baragli et al., 2011) and dingoes (Smith and Litchfield, 2010), both of which exhibit progressive improvement in obstacle navigation.

The cases of the oils containing key fatty acids, Olive oil and Flax-seed oil, demand more consideration. Fatty acids are known to play a crucial role in the interactions between ants and their environment, affecting their behaviour and ecological dynamics. Many ant species dump rubbish outside the nest in the field (e.g. *Pogonomyrmex barbatus*, Gordon, 1983). In laboratory experiments, oleic and linoleic acids have been shown to trigger removal behaviour in ants (review: Sun et al., 2018). Oleic acid, a monounsaturated fatty acid, has been tested on *Pogonomyrmex badius* and *Solenopsis saevissima* (Wilson et al., 1958), *Myrmica vindex* (Haskins and Haskins, 1974), *Myrmica rubra* (Diez et al., 2013), and *Solenopsis invicta* (Qiu et al., 2015). Linoleic acid, an omega-6 polyunsaturated fatty acid, has been tested on *Myrmica rubra* (Diez et al., 2013) and *Solenopsis invicta* (Qiu et al., 2015). In the current experiment, Olive and Flax-seed oil (sources of the aforementioned death-associated odours) showed a moderate impact on ant navigation; some of the ants detoured around while others walked over the odour line. These fatty acids had less effect here than did the strange odours of Tea tree and Lavender oils, suggesting novelty of chemical cues as a primary driver of neophobia rather than the fatty acids associated with death. Why did oils containing fatty acids shown in previous research to be associated with corpses not have a stronger effect? The context in which these fatty acids appear comes to mind as a possible reason. These food oils have all kinds of other chemicals, possibly containing odours associated with food plants. That might make the effects of oleic and linoleic acid not as strong in causing an avoidance reaction. The context of a foraging tree might also matter, as it is not a place where rubbish is found, and the ants probably have not ever encountered any dead organisms on the tree trunk during their previous foraging expeditions. In vertebrate animals (Bouton, 2004; Bouton et al., 1999) and in insects (Cheng, 2005; Colborn et al., 1999; Collett et al., 2002; Pahl et al., 2007), contextual cues modulate learning, memory, and behaviour. Further studies, however, are warranted to confirm this hypothesis, including experiments that apply pure oleic acid and linoleic acid on the foraging route, the chemicals used in the studies reviewed in this paragraph.

## CONCLUSION

In summary, this study provides important insights into the neophobic responses displayed by the bull ant *Myrmecia midas* when encountering different odours along their standard foraging routes. The results demonstrate that when exposed to unfamiliar odours, these ants exhibit a range of behaviours: avoidance of the chemicals with path deviations, scanning, and meandering movement patterns. The responses varied in intensity, with complete avoidance of foreign odours like tea tree and lavender oils, moderate avoidance of odours containing oleic and linoleic acids (which have been linked to cues associated with death), and less but still moderate avoidance for the slightly familiar eucalyptus oil. In the presence of novel odours, the ants’ paths became more curved and meandering, with reduced straightness, increased sinuosity, and lower displacement efficiency. They often decreased their speed and increased head-turning angular velocity, likely searching for alternative routes around the odour stimuli. The observed neophobic responses highlight the importance of olfactory cues in shaping the ants’ behaviour and decision making during their navigation. This study highlights the neophobia response of bull ants in response to environmental changes, showing the link between sensory perception and behavioural plasticity which enables these insects to thrive in their environments. Future research could explore the long-term effects of these odour changes on foraging efficiency and the potential that with prolonged exposure, the ants might habituate to the introduced odours.

## Acknowledgements

We acknowledge the traditional custodians of the field site on campus, the Wallumatigal clan of the Dharug Nation. The study was performed as part of a Participation and Community Engagement (undergraduate) class in biological sciences at Macquarie University. We thank Macquarie University for providing access to the field site on campus.

## Author contributions

Conception of experiments: SD; Design of experiments: VMK, YA, JL, AJ, SSN, PBC, AG, YS, SD; experimentation: VMK, YA, JL, AJ, SSN, PBC, AG, YS; data analysis: VMK, YA, JL, AJ, SSN, PBC, AG, YS, SD; supervision: SD, KC; writing of first draft: VMK, YA, JL, AJ, SSN, PBC, AG, YS; writing of revisions: KC, SD

## Ethics

Australia does not have ethical regulations concerning work with ants. The manipulations in the study were non-invasive and caused no damage to any individual ants or their colonies.

## Data availability

Supplementary videos, Excel file and R scripts are available at Open Science framework: https://osf.io/gqb8r/.

